# Genetic effects on educational attainment in Hungary

**DOI:** 10.1101/2020.01.13.905034

**Authors:** Péter P. Ujma, Nóra Eszlári, András Millinghoffer, Bence Bruncsics, Péter Petschner, Péter Antal, Bill Deakin, Gerome Breen, György Bagdy, Gabriella Juhász

## Abstract

Educational attainment is a substantially heritable trait, and it has recently been linked to specific genetic variants by genome-wide association studies (GWASs). However, the effects of such genetic variants are expected to vary across environments, including countries and historical eras. We used polygenic scores (PGSs) to assess molecular genetic effects on educational attainment in Hungary, a country in the Central Eastern European region where behavioral genetic studies are in general scarce and molecular genetic studies of educational attainment have not been previously published. We found that the PGS is significantly associated with highest educational level attained as well as the number of years in education in a sample of Hungarian volunteers (N=829). In an English (N=976) comparison sample with identical measurement protocols the same PGS had a stronger association with educational level, but not with years in education. In line with previous Estonian findings, we found higher PGS effect sizes in Hungarian, but not in English participants who attended higher education after the fall of Communism, although we lacked statistical power for this effect to reach significance. Our results provide evidence that polygenic scores for educational attainment are valid in diverse European populations.

## Introduction

Educational attainment is a key psychological and sociological variable, which comprises an important part of socioeconomic status and it is positively correlated with income and health, but negatively with crime and welfare dependency (Behrman et al., 1997). Educational attaiment is moderately heritable, with a substantial shared environmental component (Branigan et al., 2013) and it shares substantial, but not all genetic variance with cognitive abilities (Krapohl et al., 2014).

Early reports on the heritability of educational attainment were derived from family pedigree studies, most notably twin studies (Cesarini and Visscher, 2017). Recently, however, the heritability of educational attainment was confirmed with molecular genetic methods. Single nucleotide polymorphism (SNP) heritability studies (Davies et al., 2016; Hill et al., 2016) confirmed that genetic similarity between non-related individuals is positively associated with their phenotypic similarity of educational attainment, with common genotyped SNPs accounting for up to 20% of the total variance. Large-scale genome-wide association (GWA) studies linked specific genetic variants to educational attainment (Rietveld et al., 2013; Okbay et al., 2016; Lee et al., 2018). Polygenic scores based on GWAS results confirmed the predictive value of these genetic variants (also termed polygenic score [PGS] heritability), typically accounting for up to 10% of the phenotypic variance in educational attainment itself (Domingue et al., 2015; Allegrini et al., 2019), cognitive abilities (de Zeeuw et al., 2014; Selzam et al., 2016; Allegrini et al., 2019), social mobility (Ayorech et al., 2017) and overall socioeconomic success (Belsky et al., 2016; Belsky et al., 2018) in independent samples. The validity of education attaiment PGSs has been demonstrated among others in samples of Icelanders (Kong et al., 2018), Estonians (Rimfeld et al., 2018) and African Americans (Domingue et al., 2015; Lee et al., 2018; Rabinowitz et al., 2019).

However, neither the pedigree-based or SNP heritability of educational attainment nor the correlation of polygenic scores with socioeconomic phenotypes is a biological constant. There is evidence that between-country differences (Lee et al., 2018) and within-country changes in education policy (Heath et al., 1985) as well as the attendance of different types of schools (Trejo et al., 2018) may affect the heritability of educational attainment (gene-environment interaction). In other words, the relative importance of genetic and environmental effects on individual differences in educational attainment is affected by the characteristics of the environment. It has been argued (Hauser et al., 2002; Nielsen, 2008; Conley et al., 2015) that a high heritability of educational attainment is a sign of a meritocratic educational system, because attainment is determined by innate abilities and preferences instead of shared environmental effects such as social class or parental income. The social changes due to the fall of Communism (FoC) in the former Eastern Block may have had a particular effect on educational meritocracy. In line with this hypothesis, a recent Estonian study (Rimfeld et al., 2018) found that the SNP and PGS heritability of educational attainment was higher in Estonians who attended school after FoC, suggesting that the educational system in Estonia has become more meritocratic.

Because of the moderating and mediating effects of environmental variables, the strength of genetic effects on educational attainment may be different across countries. In the present study we investigated molecular genetic effects on educational attainment in Hungary, a country where no similar study has previously been published.

## Material and Methods

We used genetic data and self-reported level of education collected in the NewMood study (New Molecules in Mood Disorders, Sixth Framework Program of the European Union, LSHM-CT-2004-503474) to validate the EA3 polygenic score (Lee et al., 2018) in Hungarian participants (Budapest sample, N=829). We used data from English participants from NewMood (Manchester sample, N=976) to provide a comparison group with an identical phenotypic and genotypic data collection regimen. Participants of 18-60 years of age were recruited through advertisements, general practices and a website. Full details of the recruitment strategy and criteria have been published previously (Lazary et al., 2008; Juhasz et al., 2009; Juhasz et al., 2011). For this study the experimental cohort was limited to unrelated individuals of self-reported European white ancestry as this was the largest ethnic group. Previous studies have established the high reliability of self-reported ancestry (Tang et al., 2005; Guo et al., 2014). The study was approved by the local Ethics Committees (Scientific and Research Ethics Committee of the Medical Research Council, Budapest, Hungary; and North Manchester Local Research Ethics Committee, Manchester, UK) and was carried out in accordance with the Declaration of Helsinki and all relevant rules and regulations as part of the NewMood study. All participants provided written informed consent.

### Educational attainment

Participants filled out close-ended questions about whether they attained certain educational levels. These levels were „No qualification”, „O-levels”, „A-levels”, „Degree” „Professional qualification” and „Other (please specify)”. In the Hungarian version of the questionnaire, British educational levels were translated as their Hungarian counterparts (O-levels as „szakmunkásképző”, vocational education; A-levels as „érettségi”, high school diploma; Professional qualification as „szakvizsga”, a vocational or specialist qualification). If a participant gave a response about an „Other” qualification, it was decided individually which of the other educational levels this corresponds to based on the participant’s comments. We used these self-reported educational attainment levels to create two educational attainment phenotypes. First, we coded the highest educational level of each participant as a nominal variable (educational level). Second, we converted the self-reported educational attainment levels to years of completed education as an interval variable (years in education).

### Age groups

We aimed to investigate whether the strength of genetic effects on educational attainment varied as a function of age. In Hungary, this affected whether or not a participant graduated from high school before or after FoC, a possible moderator of the relative strength of genetic effects (Rimfeld et al., 2018). For ethical reasons, we did not store data about the birth year of our participants or exactly when they were interviewed. However, given that data collection was performed in 2004 and 2005, we can estimate the birth year of each participant within a one year margin based on self-reported age at data collection. We divided our participants in three age groups based on their age at FoC and whether they were old enough to have realized their potential for educational attainment by the time data was collected: 1) YPostC (young participants attending high school after FoC): age <24 at data collection (earliest possible birth year in 1981, possibly not old enough to have attained a university education); 2) PostC (participants attending high school mostly after FoC): age 24-31 years at data collection (birth year 1973-1981, at most 16 years old at FoC in Hungary and old enough to have attained a university education); 3) PreC (participants attending high school mostly before FoC): age at least 32 years at data collection (latest possible birth year in 1973, at least 16 years old at FoC in Hungary). We provide detailed statistics about the sample sizes, ages and educational attainments of these groups in Table 1. We hypothesized that the validity of the EA3 polygenic score will be different in these age groups in Hungary, but not in England, due to a gene-environment interaction induced by the historical political changes in Hungary and their effects on the educational system (Rimfeld et al., 2018).

**Table 1.** The distribution of age and educational level across age groups. Note that the ‘All participants’ columns under ‘Education’ also contain participants with no age data, who were consequently not assigned to either age group (N_Budapest_=54, N_Manchester_=2). For the same reason, counts in these columns are not equal to the sum of the age group columns and the total count of ‘All participants’ is different for the ‘Education’ and ‘Age’ panels.

|  | Budapest |  |  |  | Manchester |  |  |  |
| --- | --- | --- | --- | --- | --- | --- | --- | --- |
|  | YPostC | PostC | PreC | All participants | YPostC | PostC | PreC | All participants |
| EDUCATION |  |  |  |  |  |  |  |  |
| Years in education (Mean) | 12.18 | 13.68 | 13.98 | 13.25 | 14.08 | 14.11 | 13.38 | 13.71 |
| Years in education (SD) | 0.92 | 2.06 | 2.18 | 1.96 | 2.11 | 2.46 | 2.55 | 2.47 |
| No qualification | 1 | 1 | 4 | 6 | 2 | 5 | 24 | 31 |
| Professional qualification | 1 | 1 | 1 | 4 | 5 | 14 | 70 | 89 |
| O-levels or equivalent | 0 | 10 | 20 | 36 | 16 | 50 | 144 | 211 |
| A-levels or equivalent | 259 | 114 | 109 | 512 | 70 | 32 | 89 | 192 |
| University degree | 14 | 95 | 145 | 271 | 107 | 121 | 225 | 453 |
| All participants | 275 | 221 | 279 | 829 | 200 | 222 | 552 | 976 |
| **AGE** |  |  |  |  |  |  |  |  |
| 18 | 5 | 0 | 0 | 5 | 23 | 0 | 0 | 23 |
| 19 | 25 | 0 | 0 | 25 | 45 | 0 | 0 | 45 |
| 20 | 65 | 0 | 0 | 65 | 36 | 0 | 0 | 36 |
| 21 | 86 | 0 | 0 | 86 | 33 | 0 | 0 | 33 |
| 22 | 61 | 0 | 0 | 61 | 30 | 0 | 0 | 30 |
| 23 | 33 | 0 | 0 | 33 | 33 | 0 | 0 | 33 |
| 24 | 0 | 25 | 0 | 25 | 0 | 32 | 0 | 32 |
| 25 | 0 | 19 | 0 | 19 | 0 | 25 | 0 | 25 |
| 26 | 0 | 31 | 0 | 31 | 0 | 35 | 0 | 35 |
| 27 | 0 | 19 | 0 | 19 | 0 | 27 | 0 | 27 |
| 28 | 0 | 36 | 0 | 36 | 0 | 29 | 0 | 29 |
| 29 | 0 | 22 | 0 | 22 | 0 | 22 | 0 | 22 |
| 30 | 0 | 22 | 0 | 22 | 0 | 33 | 0 | 33 |
| 31 | 0 | 26 | 0 | 26 | 0 | 19 | 0 | 19 |
| 32 | 0 | 21 | 0 | 21 | 0 | 36 | 0 | 36 |
| 33-35 | 0 | 0 | 52 | 52 | 0 | 0 | 79 | 79 |
| 35-40 | 0 | 0 | 78 | 78 | 0 | 0 | 151 | 151 |
| 40-45 | 0 | 0 | 53 | 53 | 0 | 0 | 145 | 145 |
| 45-50 | 0 | 0 | 45 | 45 | 0 | 0 | 95 | 95 |
| 50-55 | 0 | 0 | 36 | 36 | 0 | 0 | 18 | 18 |
| 55-60 | 0 | 0 | 15 | 15 | 0 | 0 | 28 | 28 |
| All participants | 275 | 221 | 279 | 775 | 200 | 258 | 516 | 974 |

### Genotyping

Genomic DNA was extracted from buccal swabs collected by cytology brush (Cytobrush plus C0012; Durbin PLC). Genotyping was carried out using Illumina’s CoreExom PsychChip, genomic positions were defined according to the build GRCh37/hg19. Further details of imputation and quality control were published elsewhere (Eszlari et al., 2019).

### Statistical analysis

We used the stable 1.26.0 Genome-wide Complex Trait Analysis (GCTA) version for the calculation of SNP heritability. A minor allele frequency (MAF) cutoff of 0.05 was used. Polygenic scores were constructed using PRSice-2 and publicly available summary SNP effect size data (Lee et al., 2018), downloaded from https://www.thessgac.org/data. We used the effect sizes which were constructed without 23andMe data but released for all SNPs. A MAF threshold of 0.01 was used. In order to optimize PGS performance but avoid overfitting, we used the PGS constructed with the GWAS p-value threshold that had the strongest association with a specific phenotype, educational level within the entire sample. The best performance was achieved by PGSs with similar p-value thresholds in both samples (p=0.19 in the Budapest sample and p=0.18 in the Manchester sample), and we used these in all subsequent analyses. For nominally coded educational levels, the effect size of the association between these and PGSs was estimated as a pseudo-correlation calculated as the square root of the ratio of between-group (between-educational level) PGS variance to total PGS variance. For years in education the effect size was calculated as the Pearson correlation between PGS and this measure of educational attainment. Because the variance of years in education was not equal in all subsamples, we corrected results for restriction of range using the formula by Schmidt & Hunter (Schmidt and Hunter, 2014).

We ran both GCTA and PGS analyses both with and without controlling for the first 10 genomic principal components.

### Availability of data

The data that support the findings of this study are available on request from the corresponding author. The data are not publicly available due to privacy or ethical restrictions.

## Results

Genomic-relationship-matrix restricted maximum likelihood (GREML-GCTA) SNP heritabilities indicated that in the Budapest sample, all common SNPs accounted for 30.5% (SE=25%, p=0.09) of the variance of educational level and 34.4% (SE=24%, p=0.06) of the variance of years in education. In the Manchester sample, the same SNP heritabilities were 58% (SE=20%, p=0.01) for educational level and 20.5% (SE=20%, p=0.13) for years in education. Controlling for the first 10 genomic PCs, the corresponding values were h^2^_SNP_=34.3% (Budapest, educational level, SE=25.3%, p=0.07), h^2^_SNP_=42.6% (Budapest, years in education, SE=24.6%, p=0.03), h^2^_SNP_=58.2% (Manchester, educational level, SE=21.1%, p=0.001), h^2^_SNP_=20.2% (Manchester, years in education, SE=20.7%, p=0.147). These estimates were likely biased downward due to low variance in the youngest participants, many of whom were still in education (Table 1). However, as our sample sizes were below the several thousand individuals usually recommended for this type of analysis (Knopik et al., 2016) and because GCTA models failed to properly converge when we further restricted samples to single age groups due to very low sample sizes we did not perform SNP heritability analyses within these separately. The EA3 PGS was significantly associated with both educational attainment phenotypes in both the total Budapest (R_education level_=0.14, r_years, corrected_=0.16) and Manchester (R_education level_=0.3, r_years, corrected_=0.22) samples. Multiple regressions adding age, sex and the first 10 genomic PCs as covariates improved model fit (Supplementary table S1), but did not meaningfully change the relationship between PGS and educational attainment (absolute Δβ_max_=0.018, average Δβ=−0.003).

Figure 1 shows group means and the dispersion of PGSs by country, age group and educational attainment level. The highest mean values were always found in university-educated participants. High school-educated participants had lower means, followed by individuals with vocational educations (O-levels in the Manchester sample). Individuals with professional educations had somewhat higher means. We note, however, that the low number of participants with low educational attainments led to less precision in estimating mean PGSs. Note especially the limited educational attainment variability in the Budapest sample, with most participants having either high school or university educations.

**Figure 1.**
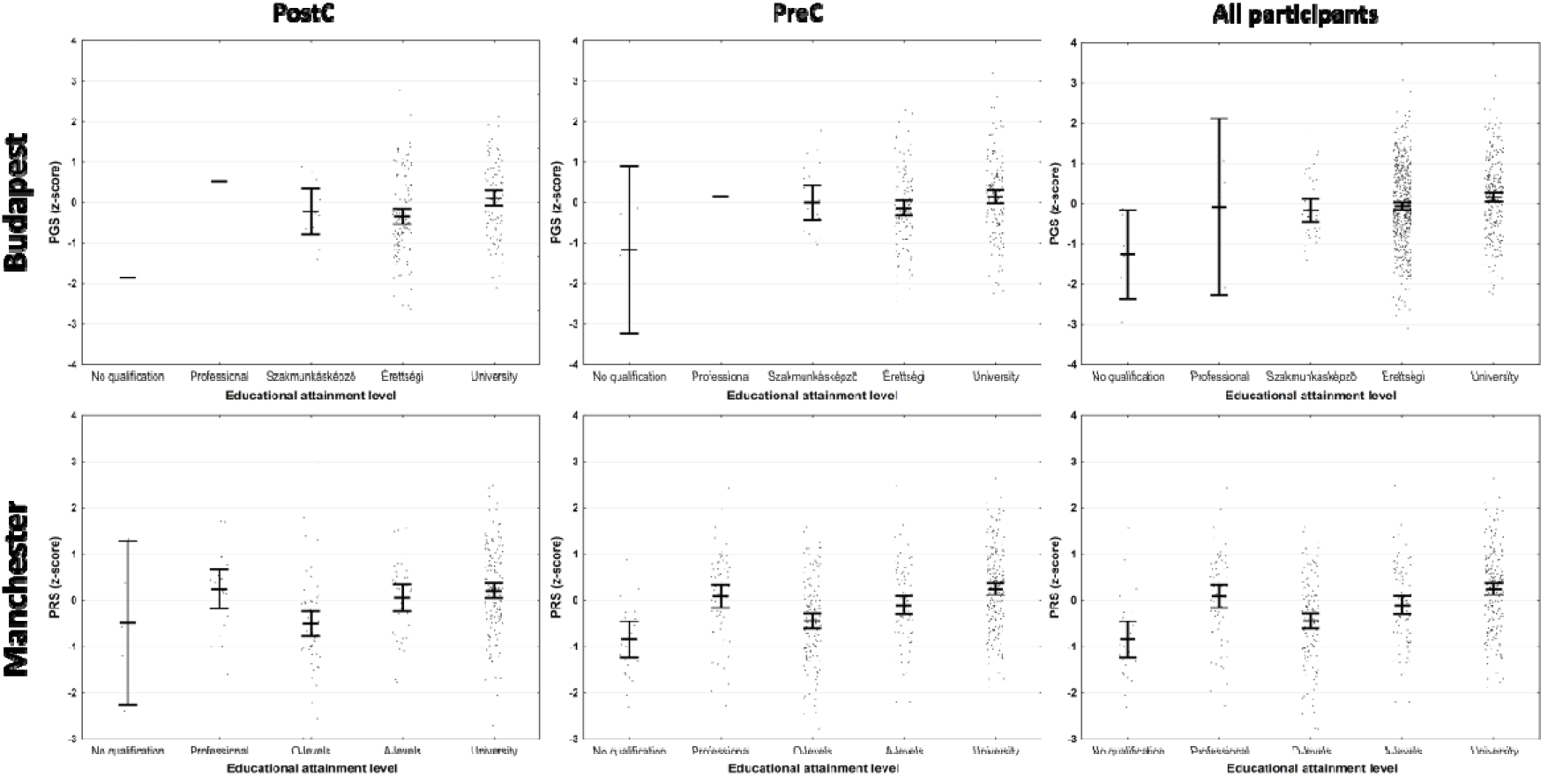
PGSs by country and educational levels. PostC and PreC indicate age groups, see Table 1. „All participants” includes partipants with no age data. Educational attainment categories are equivalent, but indicated with the original questionnaire terms for authenticity. In the Budapest sample, „Szakmunkásképző” refers to a vocationally-oriented high school education, while „Érettségi” refers to high school education with a successful final standardized examination. Central horizontal lines show group means and whiskers indicate 95% confidence intervals (CIs), overplotted with raw data. Note that some Budapest groups were represented by a single participant which did not permit the estimation of CIs.

We also calculated PGS-phenotype associations by country and age group separately (Figure 2), except the youngest participants (age<24) from these analyses because the low variability of educational attainment in this subgroup (see Table 1). We also report findings from the pooled PreC and PostC subsamples (that is, from all participants whose age was known and at least equal to 24 years), which are less affected by individuals likely still in education. The PGS was significantly associated with both phenotypes in all individual subsamples (effect size=0.13-0.34).

**Figure 2.**
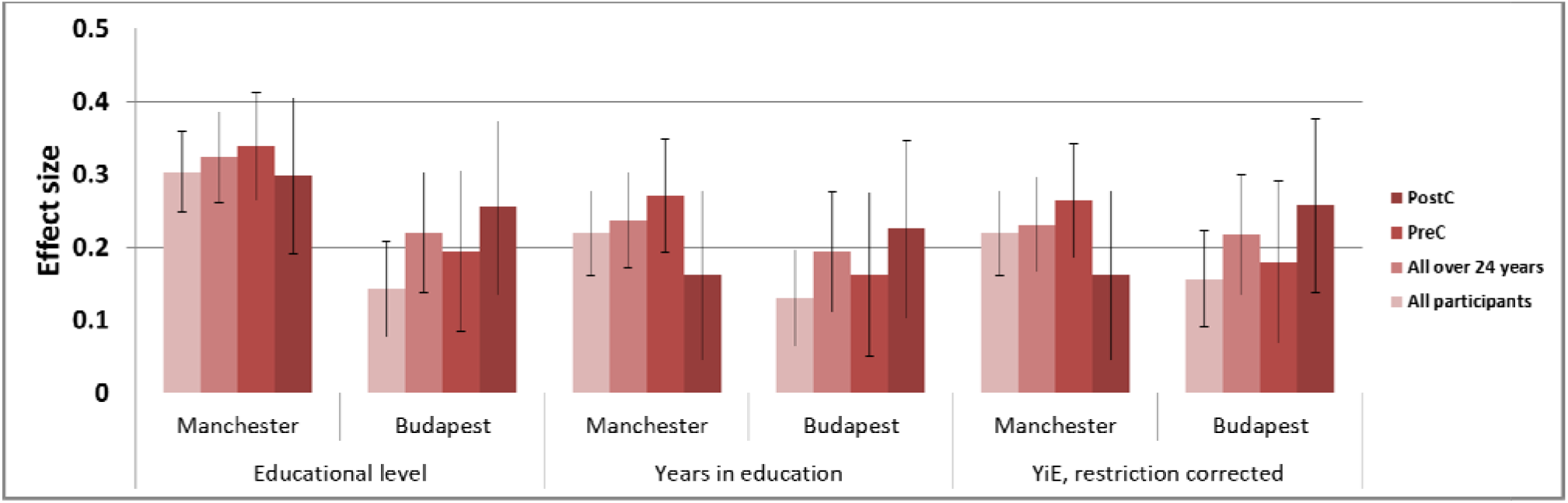
Associations between the best-fit EA3 polygenic score and educational attainment by sample and age group. For the nominally coded educational level, the effect size is a pseudo-correlation equal to the the square root of the proportion of between-levels PGS variance to total PGS variance. For the years in education, the effect size is a Pearson correlation coefficient. Error bars show 95% CIs. „YiE, restriction corrected” refers to a Pearson correlation coefficient between the PGS and years in education, corrected for restriction of range. PostC: participants at most 16 years old at FoC and at least 24 years old during data collection. PreC: participants at least 16 years old at FoC. „All over 24 years” refers to pooled PostC and PreC subsamples. „All participants” also includes participants younger than 24 years old at data collection and those with no age data.

In order to assess age group and country effects on the relative strength of PGS effects, we compared effect sizes of each phenotype (educational level, years in education, years in education corrected for restriction of range) between all subgroups using Fisher’s r-to-z method (Rimfeld et al., 2018). Effect sizes were significantly higher in the Manchester sample for educational level, but not for years in education, especially after restriction of range was corrected for. In line with previous Estonian results, PostC Budapest subsamples had 25-30% higher effect sizes than the PreC subsamples for all phenotypes, but this did not reach statistical significance at our sample size (p_min_=0.37). We report detailed results in Supplementary table S2. We set the age cutoff between PreC and PostC groups at 32 years in 2004-2005 (16-17 years old at FoC) following Rimfeld et al, but still somewhat arbitrarily. In order to test the effect of different age cutoffs on the results, we performed a specification curve analysis using all possible PreC/PostC age cutoffs between 26-45 years in 2004-2005 (Supplementary figure S1). In line with previous analyses, we never included those under 24 years old in either the PreC or the PostC groups. For both educational level and years in education, the trend of higher PGS effect sizes in the Budapest PostC sample persisted for all cutoffs less than 40 years (birth year: 1964/65, at least 25 years old at FoC) with no similar effect in the Manchester sample. However, at this sample size no age cutoff yielded a statistically significant difference between the effect sizes in the PreC and PostC subsamples for either phenotype and sample.

## Discussion

Ours is the first study to estimate molecular genetic effects on educational attainment in Hungary, and the second to do so in a former Warsaw Pact country. We are also unaware of any other behavior genetic study about educational attainment or cognitive functions in Hungarians. We found that – in line with international results – a substantial proportion of educational attainment variance is accounted for by common genetic variants in Hungarians, and that the genetic variants discovered by a recent GWAS to predict educational attainment in Western European and American validation samples also do so in Hungary.

We compared findings in Hungarians to analogous results from an English comparison sample with identical recruitment protocols and phenotypic measurements. Once restriction of range was corrected for either statistically or by excluding very young participants presumably still in education, PGS effect sizes were either not significantly lower in the Budapest sample (for years in education) or they only reached marginal significance in the comparison of specific subsamples (for educational level). We note that there was much higher variability in educational level in the Manchester sample (Table 1) which likely contributed to the effect size differences between samples, but a formal correction of restriction of range was not possible for this nominally coded variable.

An exact comparison of our PGS effect sizes with other studies is not feasible due to between-study differences in genotyping, polygenic score construction (such as differences in the source GWAS and the selection of p-value and MAF thresholds) and phenotype quality (including the specific phenotype used and its variance). However, we note that that the effect sizes in the Manchester sample were in line with those reported by indepedent studies using PGSs based on the same GWAS with more representative British and American datasets with higher quality phenotypes (Lee et al., 2018; Allegrini et al., 2019) including educational attainment and cognitive performance. The effect sizes in the Budapest sample were generally not substantially weaker than this. In sum, the relative strength of genetic effects in our Budapest sample were in line with those reported from other countries.

Our aim to replicate previous Estonian findings about the larger relative role of genetic effects after FoC was only partially successful. While we found substantially higher effect sizes for educational attainment phenotypes in the FoC subsamples in the Budapest, but not the Manchester sample, these differences did not reach statistical significance. Changes in educational policy surrounding FoC were similar in Hungary to Estonia (Ladányi, 1995; Hrubos et al., 2016). The previous Estonian study on the same effects (Rimfeld et al., 2018) invokes increases in educational meritocracy – first of all, the abandonment of political considerations in university admissions – as the chief driver of increased PGS effect sizes after FoC. Our results do not exclude the possibility of a similar change taking place in Hungary, but better powered genetic databases will be required for a conclusive replication.

Our work suffers from a number of limitations. The largest of these is the modest size of our sample, which allowed us to conclusively demonstrate the association of the PGS polygenic score with actual educational attainment in Hungarians, but limited statistical power to detect age and country effects on SNP heritability (Supplementary Table S2). Systematically higher PGS effect sizes in the FoC Budapest sample suggest that a historical gene-environment interaction may have taken place in Hungary during FoC, but this requires replication in larger samples. We note, however, that large, approximately population-representative genetic databases like the Estonian Biobank are rare in Central and Eastern Europe, and therefore ours is probably the best Hungarian dataset currently available to test our hypotheses. Second, our database was not nationally representative and most educational attainment differences existed between completion and non-completion of college. This may have exerted a downward bias on our PGS validity estimates due to range restriction, especially in the Hungarian sample. We corrected our results for restriction of range, but only relative to the subsample with the highest variance. Furthermore, we are unaware of any statistical method which can correct for restriction of range in a nominal variable, and therefore did not attempt such a correction for educational level.

Third, a general limitation to between-family molecular genetic studies is that they may reveal shared environmental instead of true, biological genetic effects through gene-environment correlation (Young, 2019). One the one hand, SNPs used to construct PGSs or the relatedness matrix for GCTA may be associated with eduational attainment because they index membership in families which influence educational attainment through cultural rather than genetic effects (residual stratification or ‘dynastic’ effects (Morris et al., 2019)). On the other hand, SNPs may have a causal effect on parental phenotypes which in turn influence offspring educational attainment (genetic nurture (Kong et al., 2018)). While both effects are known to operate and inflate PGS effect sizes in studies of unrelated individuals (Kong et al., 2018; Young et al., 2018; Young, 2019), validity in within-family studies (Domingue et al., 2015; Belsky et al., 2018; Selzam et al., 2019) demonstrates that a substantial portion of PGS effects reflects actual genetic influences. While we do not expect the effect of genetic nurture to be substantially different in Hungary than in other countries, we expect residual stratification to inflate PGS effect sizes more in our Manchester sample and other British between-family studies than in our Budapest sample. This is because the GWAS based on which we constructed our PGSs contained substantial British study populations, but no Hungarians and a very limited number of participants from neighboring countries (specifically, 777 Austrians and 842 Croatians) which would bias stratification effects towards patterns which exist in Britain. Future Hungarian within-family studies may provide a formal test of this hypothesis.

Finally, the investigated after-FoC period ends in 2004-2005 at the time of data collection and our study does not investigate educational attainment after this time.

In sum, our work demonstrates that genetic variants discovered in international GWAS samples also predict educational attainment in Hungary with equal or only slightly reduced strength relative to an English sample. In line with Estonian data, individual genetic differences played a larger role shaping educational attainment in those graduating after the fall of Communism, but due to limitations in statistical power a more conclusive replication of this effect is needed. Similar findings from Hungary had not been previously available, and the results are likely of interest to those studying the society of Hungary and may serve as a model for other countries of the region without their own genetic studies.

## Acknowledgements

The study was supported by the Sixth Framework Program of the European Union (NewMood, LSHM-CT-2004-503474), by the Hungarian Brain Research Program (Grants: 2017-1.2.1-NKP-2017-00002 and KTIA_13_NAPA-II/14), the National Development Agency (Grant: KTIA_NAP_13-1-2013-0001), the Hungarian Academy of Sciences, Hungarian National Development Agency, Semmelweis University and the Hungarian Brain Research Program (Grant: KTIA_NAP_13-2-2015-0001, MTA-SE-NAP B Genetic Brain Imaging Migraine Research Group), and by the Hungarian Academy of Sciences (MTA-SE Neuropsychopharmacology and Neurochemistry Research Group). It was also supported by the National Institute for Health Research Manchester Biomedical Research Centre, by TAMOP-4.2.1.B-09/1/KMR-2010-0001, by OTKA 119866, by the BME-Biotechnology FIKP grant of EMMI (BME FIKP-BIO), by the New National Excellence Program of The Ministry of Human Capacities (ÚNKP-16-3, ÚNKP-17-3-III-SE-2, and ÚNKP-17-4-I-SE-8), by ITM/NKFIH Thematic Excellence Programme, Semmelweis University; and by the SE-Neurology FIKP grant of EMMI.

