## Supplementary Table S1 for "Genetic effects on educational attainment in Hungary"

| **Budapest** |  | **Model 1** | | |  | **Model 2** | | | **Model 3** | | | | |  |  |
| --- | --- | --- | --- | --- | --- | --- | --- | --- | --- | --- | --- | --- | --- | --- | --- |
|  |  | **β** | **t** | **p** | **Model R^2^** | **β** | **t** | **p** | **Model R^2^** | **β** | **t** | **p** | **Model R^2^** | **Δβ_1-2_** | **Δβ_1-3_** |
| PreC | PGS | 0.163 | 2.745 | 0.006 | 0.026 | 0.167 | 2.805 | 0.005 | 0.032 | 0.155 | 2.517 | 0.012 | 0.047 | 0.004 | -0.008 |
|  | Sex |  |  |  |  |  | -0.153 | 0.879 |  |  | -0.109 | 0.891 |  |  |  |
|  | Age |  |  |  |  | -0.075 | -1.260 | 0.209 |  | -0.062 | -0.884 | 0.321 |  |  |  |
| PostC | PGS | 0.225 | 3.414 | <0.001 | 0.051 | 0.232 | 3.526 | <0.001 | 0.065 | 0.238 | 3.481 | <0.001 | 0.125 | 0.008 | 0.013 |
|  | Sex |  |  |  |  |  | -0.712 | 0.483 |  |  | -0.655 | 0.513 |  |  |  |
|  | Age |  |  |  |  | 0.115 | 1.752 | 0.081 |  | 0.110 | 3.481 | 0.102 |  |  |  |
| All over 24 years | PGS | 0.194 | 4.401 | <0.001 | 0.037 | 0.193 | 4.365 | <0.001 | 0.039 | 0.199 | 4.392 | <0.001 | 0.059 | <0.001 | 0.005 |
|  | Sex |  |  |  |  | 0.021 | -0.524 | 0.600 |  |  | -0.448 | 0.654 |  |  |  |
|  | Age |  |  |  |  | -0.023 | 0.481 | 0.630 |  | 0.026 | 0.576 | 0.565 |  |  |  |
| All participants | PGS | 0.127 | 3.760 | <0.001 | 0.017 | 0.117 | 3.437 | <0.001 | 0.113 | 0.109 | 3.543 | <0.001 | 0.128 | -0.010 | -0.018 |
|  | Sex |  |  |  |  | 0.050 | 1.609 | 0.259 |  |  | 1.570 | 0.117 |  |  |  |
|  | Age |  |  |  |  | 0.301 | 8.915 | <0.001 |  | 0.303 | 8.863 | <0.001 |  |  |  |
| **Manchester** |  |  | | |  |  | | |  |  |  |  |  |  |  |
|  |  | **β** | **t** | **p** | **Model R^2^** | **β** | **t** | **p** | **Model R^2^** | **β** | **t** | **p** | **Model R^2^** | **Δβ_1-2_** | **Δβ_1-3_** |
| PreC | PGS | 0.271 | 6.386 | <0.001 | 0.026 | 0.272 | 6.496 | <0.001 | 0.026 | 0.264 | 6.21 | <0.001 | 0.044 | 0.001 | -0.007 |
|  | Sex |  |  |  |  |  | 1.799 | 0.073 |  |  | 1.733 | 0.084 |  |  |  |
|  | Age |  |  |  |  | -0.171 | -4.077 | <0.001 |  | -0.178 | -4.228 | <0.001 |  |  |  |
| PostC | PGS | 0.161 | 2.613 | 0.010 | 0.074 | 0.161 | 2.591 | 0.010 | 0.106 | 0.165 | 2.591 | 0.010 | 0.128 | -0.001 | 0.004 |
|  | Sex |  |  |  |  |  | 0.297 | 0.767 |  |  | 0.287 | 0.774 |  |  |  |
|  | Age |  |  |  |  | 0.003 | 0.045 | 0.964 |  | -0.012 | -0.195 | 0.845 |  |  |  |
| All over 24 years | PGS | 0.238 | 6.809 | 0.000 | 0.057 | 0.232 | 6.748 | <0.001 | 0.092 | 0.227 | 6.554 | <0.001 | 0.104 | -0.006 | -0.001 |
|  | Sex |  |  |  |  |  | 1.621 | 0.105 |  |  | 1.597 | 0.111 |  |  |  |
|  | Age |  |  |  |  | -0.184 | -5.350 | <0.001 |  | -0.184 | -5.339 | <0.001 |  |  |  |
| All participants | PGS | 0.219 | 7.025 | <0.001 | 0.048 | 0.206 | 6.653 | <0.001 | 0.075 | 0.203 | 6.46 | <0.001 | 0.085 | -0.013 | -0.016 |
|  | Sex |  |  |  |  |  | 1.825 | 0.068 |  |  | 1.824 | 0.068 |  |  |  |
|  | Age |  |  |  |  | -0.161 | -5.204 | <0.001 |  | -0.160 | -5.145 | <0.001 |  |  |  |

Supplementary Table S1. Univariate ANOVA results with Years in education as the dependent variable and the polygenic score (Model 1), the polygenic score, sex and age (Model 2) or the polygenic score, age, sex and the first 10 genomic principal components (Model 3) as independent variables. Beta coefficients in Model 1 are identical to zero-order Pearson correlation coefficients. Δβ_1-2_ and Δβ_1-3_ indicates the change of the PGS beta coefficient from Model 1 to Model 2 and from Model 1 to Model 3, respectively.

|  | **Educational level** |  |  |  |  |  |  |  |  |
| --- | --- | --- | --- | --- | --- | --- | --- | --- | --- |
|  |  | **BP_PostC** | **BP_pre** | **BP_all_24** | **BP_all** | **MAN_PostC** | **MAN-pre** | **MAN_all_24** | **MAN_all** |
|  | **BP_PostC** |  | 0.695 | 0.443 | 1.536 | 0.512 | 1.143 | 0.980 | 0.713 |
|  | **BP_pre** | 0.487 |  | 0.359 | 0.777 | 1.268 | **2.081** | **1.969** | 1.707 |
|  | **BP_all_24** | 0.658 | 0.720 |  | 1.426 | 1.080 | **2.040** | 1.933 | 1.622 |
|  | **BP_all** | 0.125 | 0.437 | 0.154 |  | **2.291** | **3.724*** | **3.836*** | **3.601*** |
|  | **MAN_PostC** | 0.609 | 0.205 | 0.280 | **0.022** |  | 0.590 | 0.387 | 0.089 |
|  | **MAN-pre** | 0.253 | **0.037** | **0.041** | **<0.001*** | 0.555 |  | 0.302 | 0.714 |
|  | **MAN_all_24** | 0.327 | **0.049** | 0.053 | **<0.001*** | 0.698 | 0.763 |  | 0.451 |
|  | **MAN_all** | 0.476 | 0.088 | 0.105 | **<0.001*** | 0.929 | 0.476 | 0.652 |  |
| **Years in education** | |  |  |  |  |  |  |  |  |
|  |  | **BP_PostC** | **BP_pre** | **BP_all_24** | **BP_all** | **MAN_PostC** | **MAN-pre** | **MAN_all_24** | **MAN_all** |
|  | **BP_PostC** |  | 0.712 | 0.403 | 1.292 | 0.716 | 0.610 | 0.172 | 0.076 |
|  | **BP_pre** | 0.476 |  | 0.424 | 0.487 | 0.018 | 1.526 | 1.108 | 0.863 |
|  | **BP_all_24** | 0.687 | 0.672 |  | 1.156 | 0.433 | 1.304 | 0.798 | 0.491 |
|  | **BP_all** | 0.196 | 0.627 | 0.248 |  | 0.451 | **2.628** | **2.227** | 1.959 |
|  | **MAN_PostC** | 0.474 | 0.986 | 0.665 | 0.652 |  | 1.507 | 1.097 | 0.859 |
|  | **MAN-pre** | 0.542 | 0.127 | 0.192 | **0.009** | 0.132 |  | 0.635 | 1.008 |
|  | **MAN_all_24** | 0.864 | 0.268 | 0.425 | **0.026** | 0.273 | 0.525 |  | 0.391 |
|  | **MAN_all** | 0.940 | 0.388 | 0.624 | 0.050 | 0.391 | 0.313 | 0.696 |  |
| **Years in education, corrected for restriction of range** | | | | | |  |  |  |  |
|  |  | **BP_PostC** | **BP_pre** | **BP_all_24** | **BP_all** | **MAN_PostC** | **MAN-pre** | **MAN_all_24** | **MAN_all** |
|  | **BP_PostC** |  | 0.893 | 0.523 | 1.375 | 1.086 | 0.095 | 0.354 | 0.534 |
|  | **BP_pre** | 0.372 |  | 0.513 | 0.341 | 0.221 | 1.187 | 0.767 | 0.601 |
|  | **BP_all_24** | 0.601 | 0.608 |  | 1.096 | 0.749 | 0.797 | 0.266 | 0.045 |
|  | **BP_all** | 0.169 | 0.733 | 0.273 |  | 0.064 | **1.999** | 1.548 | 1.367 |
|  | **MAN_PostC** | 0.278 | 0.825 | 0.454 | 0.949 |  | 1.407 | 1.010 | 0.855 |
|  | **MAN-pre** | 0.924 | 0.235 | 0.426 | **0.046** | 0.159 |  | 0.611 | 0.874 |
|  | **MAN_all_24** | 0.723 | 0.443 | 0.790 | 0.122 | 0.312 | 0.541 |  | 0.266 |
|  | **MAN_all** | 0.594 | 0.548 | 0.964 | 0.171 | 0.393 | 0.382 | 0.790 |  |

Supplementary Table S2. Age and country differences in the PGS effect sizes. Identical tables are reported for all three phenotypes (educational level, years in education, years in education

corrected for restriction of range). The upper diagonals contains z-values and the lower diagonal contains p-values for the relevant comparisons (bold denotes nominal significance at p-value*<0.05*, an additional asterisk denotes significance after correcting for multiple comparisons using the FDR method (Benjamini and Hochberg, 1995)). BP: Budapest sample, MAN: Manchester sample. PostC and PreC indicate age groups, see Table 1. „All” indicates all participants, including those with no age data.

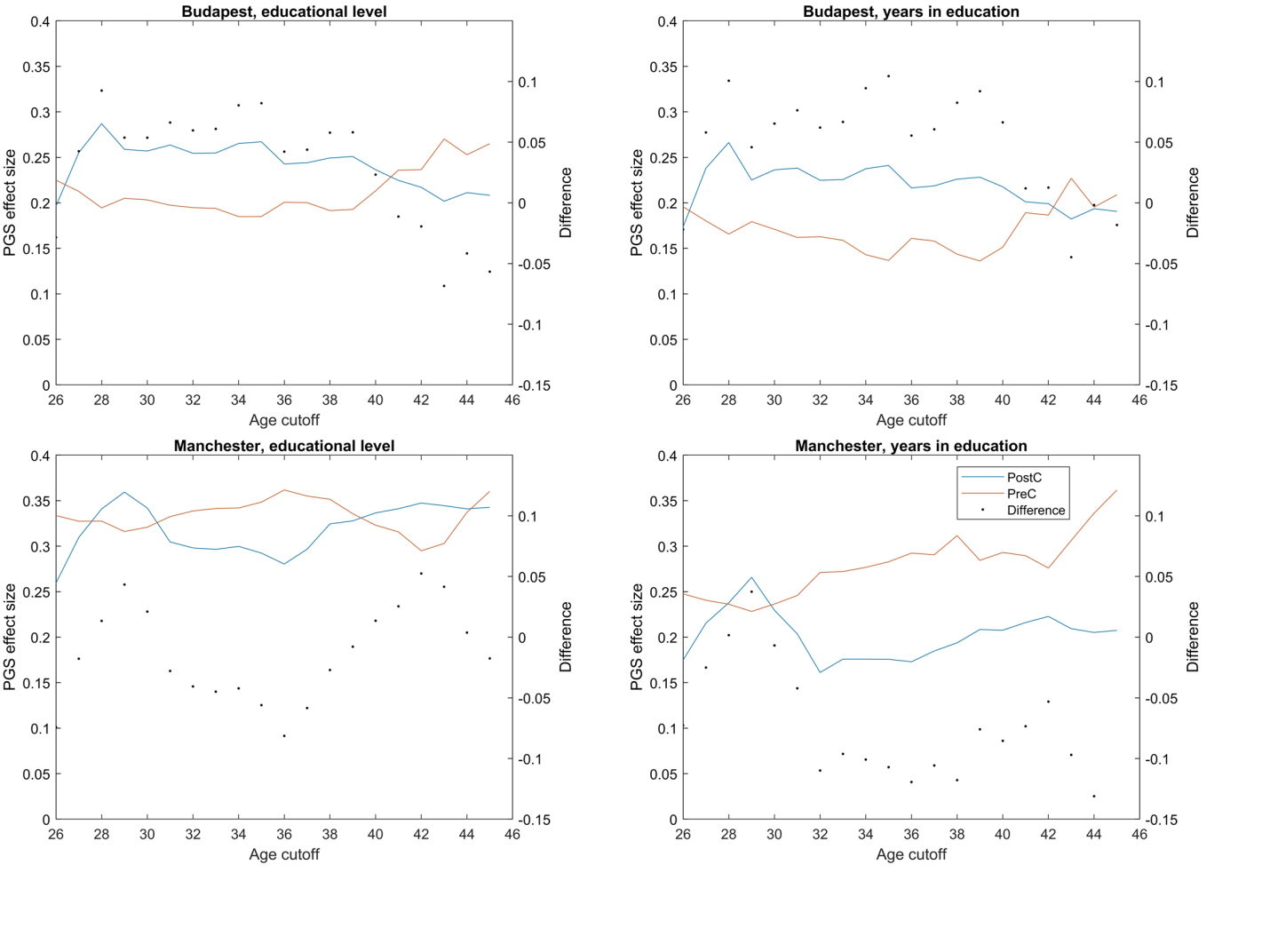

Supplementary Figure S1. Specification curve analysis of the effect of age cutoff choice on PreC-PostC PGS effect size differences. The lines indicate PostC and PreC effect sizes as a function of the age cutoff used to construct these categories (originally: 32 years) while dots indicate differences, shown on the right y axis. No PreC-PostC effect size pair differences reach statistical significance.
